# Playback-aided surveys and acoustic monitoring in the detection of the Endangered Forest Owlet *Athene blewitti*

**DOI:** 10.1101/2024.05.05.592555

**Authors:** Amrutha Rajan, Aditi Neema, Pranav G Trivedi, Sejal Worah, Meera MR, Shomita Mukherjee, V.V. Robin

## Abstract

Long-term monitoring of populations of rare, endangered species is often challenging. Both the availability of baseline historical datasets and appropriate methods for long-term monitoring are often limited. Anthropogenic climate change and landscape change can impact species distributions significantly, sometimes resulting in a distributional shift of species and, in some cases, driving species to extinction. The Forest Owlet is an endangered bird that was considered extinct but was rediscovered after 113 years in 1997. Since its rediscovery, followed by the description of its calls, there have been regular sightings of the species from newer locations, leading to its down-listing in the IUCN Red List from Critically Endangered to Endangered. One area of interest is the Dang region in Gujarat, India, where there have been no historical records despite previous ornithological studies, but there are several recent records. Through field surveys, we repurpose data from the last three decades (1990, 2000, and 2019) to explore if this bird currently occurs in previous study sites where it was not recorded. The period has seen the rise of acoustic data, and we assess if new survey techniques using playback of its call could increase its detection. Additionally, we examined any changes in landscape and climate in this region across the same period. We also developed an acoustic detection framework for detecting the Forest Owlet from co-occurring sympatric owlets using Automated Recording Units (ARU) and sound-analysis software. We assessed appropriate detection distances from vocalizing birds on the field to design a spacing grid for future surveys of the species. We could locate the Forest Owlet from the resurveys at locations where previous studies three decades ago had not. We also find a significant change in the landscape - loss of native forests and increased agriculture, along with a significant change in climatic variables - mean maximum temperature and mean rainfall. Although the detection of the Forest Owlet is higher when accompanied by playback of its call, there is considerable variation across the landscape. Our acoustic detector comparison led us to a detection strategy for long-term monitoring – different approaches for songs and calls, and an effective detection distance of 300m in its habitat. Although the species responds to climate and habitat change, our study cannot determine the cause of the increased reports of this endangered species. All possibilities remain; the increase in the recent records of the species could be from variable detection or changes in climate and land use. However, we do find increased detections with newer survey techniques with bioacoustics, and we recommend they be used with care for future baseline studies that are urgently required.

## INTRODUCTION

Anthropogenic climate change and land-use changes are known to have significantly affected biodiversity (Blake and Loiselle 2015; Mantyka-Pringle et al. 2015; Skogen, Helland, and Kaltenborn 2018). We are limited in our ability to detect such changes in biodiversity due to the lack of appropriate comparative datasets, or challenges with detecting cryptic species. Many cryptic species escape detection through traditional surveys for lack of appropriate search images (Lauriault and Wiersma 2019), and in modern times are enhanced through the use of camera traps (Thomas et al. 2020), and Acoustic Recording Units (ARU) (Frommolt and Tauchert 2014; Zwart et al. 2014; Bobay, Taillie, and Moorman 2018) deployed over longer periods to increase the chances of detecting such species.

The best-known examples of recent resurvey of avian diversity are the comparisons with the early 20th-century Joseph Grinnell surveys indicating a loss of up to 43% of species (Iknayan and Beissinger 2018; Morgan W. Tingley et al. 2012; M. W. Tingley and Beissinger 2013). They also indicated that the Mojave bird community collapse was affected by climate change, chiefly through precipitation changes rather than temperature. A key element of any resurvey is the availability of systematic historical data that can be adapted to modern methods of biodiversity surveys. From the biodiversity hotspots of India, there are very few such datasets and even fewer resurveys like Sashikumar et al. (2014) and Subramanya (2019).

The Forest Owlet *Athene blewitti* is endemic to India, occurring only in small parts of central India and the northern Western Ghats (BirdLife International 2018). After 113 years, it was rediscovered in 1997 in north Maharashtra (King and Rasmussen 1998). Since its rediscovery, new records have emerged at various locations in Madhya Pradesh, Maharashtra, and Gujarat (Mehta et al. 2017). Until 2017, its population was assessed to be less than 250 individuals remaining in the wild, and was listed as a Critically Endangered species (BirdLife International 2016). However, the population is now estimated to be up to a thousand individuals, and the species was down-listed as an endangered species in 2017 (BirdLife International 2018).

Most significantly, the first recorded call and three types of vocalizations of the species were described recently (Rasmussen and Ishtiaq 1999; Ishtiaq and Rahmani 2004). The location records of this species have increased since the recordings of its vocalisations have become more accessible. Today, playback of its calls is used in various detection-based studies (Mehta et al. 2008, 2017; Khan et al. 2023)

One of the areas where the Forest Owlet is being increasingly recorded is the Dang area in Gujarat, at the northern edge of the Western Ghats. The first records of Forest Owlet from Dang were in 2014 (Patel et al. 2015), and since then, they have been increasingly recorded from this region. Although Salim Ali’s landscape-wide bird surveys in 1954 (Ali 1954) covered the Dang, Forest Owlet was not detected here, and neither have ornithological studies in 1990 (Worah 1991) and 2000 (Trivedi 2006). It is unclear if these birds have recently moved into this landscape or if the recent targeted surveys with different methods have enhanced the detection of the species.

Although it may not be possible to answer this question comprehensively, we repurposed two previous datasets of surveys from 1990 (Worah 1991), 2000 (Trivedi 2006), and a recent occupancy survey in 2019 (Khan et al. 2023) and re-surveyed these locations with traditional and acoustic-aided methods. We also examined differences in climate in the time intervening the surveys. In parallel, we assessed the efficiency of purely acoustics-based detection methods for the long-term and targeted monitoring of the species.

The development of Automated Recording Units (ARU) and their associated algorithms have enabled researchers to conduct long bioacoustic deployments. The clear advantage of using bioacoustic sensors is obtaining long-deployment data with vocalizations of rare species that may be difficult to detect with infrequent human visits. However, one of the challenges that remain with using ARU-generated bioacoustic data is the validation of the presence of a particular species and the assessment of uncertainty around the detection of that species. Investigators have shown that background noise, the presence of other sonically similar species can obfuscate the presence of a target species (Ralph et al. 1993). The detection distance of a species in a specific habitat can impact detection and needs to be ascertained for each study (Piercy et al. 2014; Haupert, Sèbe, and Sueur 2023). Previously, a study showed the Forest Owlet call can be heard at a distance of 1 km (Ishtiaq and Rahmani 2004); however, this species has a variety of vocalisations that have yet to be assessed systematically. ARUs have been used to detect several endangered species (Arvind et al. 2023), and this is proving to be an efficient tool for the rapid assessment of threatened taxa.

In this study, we use a cascading set of questions to examine the changes in Forest Owlet detections over the years at a location - Dang Forests in Gujarat, while exploring different detection methods for future surveys.

First, we ask a) Do Forest Owlet currently occur at sites that were previously surveyed thrice in the past (1990 to 2019)

Next, we ask b) If detections of the Forest Owlet at the resurvey locations are similar with playback (newer methods) and without (traditional survey methods).

We also ask c) If the landscape or climate has changed in the period between the surveys.

Based on the role of acoustics in detections, we further assessed the efficacy of using bioacoustics-based detection of the Forest Owlet by asking:Which acoustic analysis software performs better at detecting the calls and songs of the Forest Owlet with co-occurring owls and e) At what distance should Automated Recording Units (ARU) be placed to record the species in its habitat?

## METHODS

This study was conducted in the northernmost part of the Western Ghats - Dang district of Gujarat (20°33’40", 21°5’10" North and 73°27’58", 73°56’36" East) (Figure 1), where a large portion (1035 sq. km) of the 1764 sq. km. is forested, and is spread across elevations ranging between 300 to 700 MSL.

**Figure 1:**
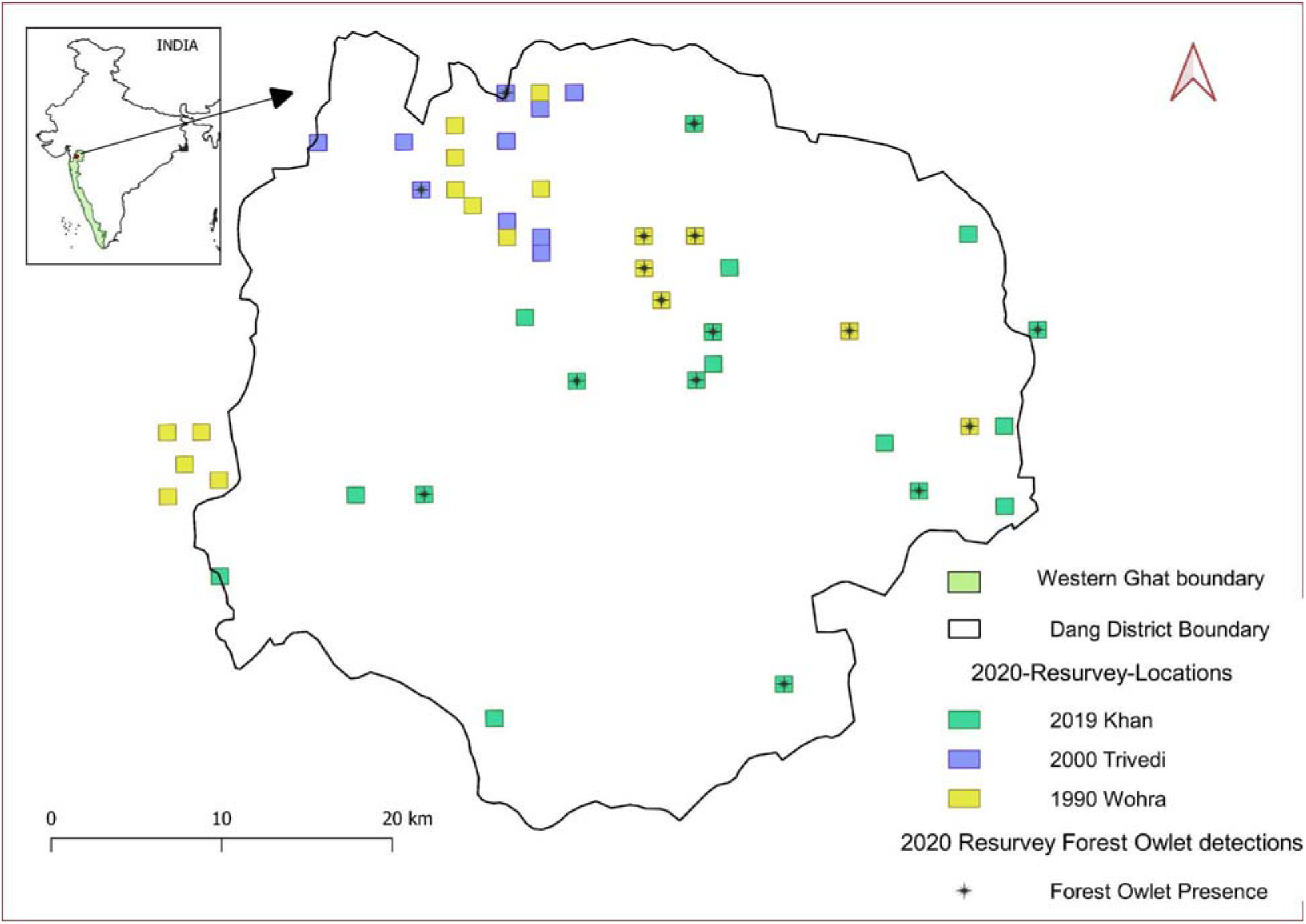
Resurveys for the endangered Forest Owlet in the Dang region result in detections at locations where studies three decades back did not

### **a)** Do Forest Owlets currently occur at sites that were previously surveyed?

We collated three past datasets across which we based our resurveys. The earliest dataset was from 1988-1991(Worah 1991), denoted hereafter as Worah 1990 by the year of the survey; and a decade later, in 2000 (Trivedi 2006), hereafter Trivedi 2000; and the latest, almost two decades later in 2018-2019 by (Khan et al. 2023) - hereafter Khan 2019. All study years are denoted by the year of surveys and not by the year of publication.

To assess any changes in climate and landscape, along with changes in the presence of the Forest Owlet, we also collated secondary datasets available for this region. Since climate and landscape change can have effects at a larger scale than the immediate location of occurrence, we included a buffer of 5 km around the Dang region - thus estimating the change in climate and landscape over 8245 sq. km.

### Historical datasets assessed

***i) Worah 1990*** was a study of the entire bird community, without a specific search image for the Forest Owlet since it was not recorded from this region at that time. Worah used 5-minute counts of all birds that were seen or heard across 74 sq. km. of the landscape over three years (1988-1991).
***ii) Trivedi 2000*** used ten line transects (1 to 3.5 km length, open width) across ∼160.84 sq km to conduct bird counts through the year for two years (2001-2003) in the same landscape. During this survey too, Forest Owlets were not recorded from the landscape, and no particular effort was made to look for the species.
***iii) Khan 2019*** is not a historical dataset, but these surveys were conducted between November 2018 to January 2019, specifically looking for Forest Owlet with active playbacks of the species. They surveyed 45 randomly selected grids across the Dang area, covering 2.5% of the total area. These surveys targeted detecting the Forest Owlet, and the team played back calls of Forest Owlet for 5 minutes (or until detection if that was earlier). They walked through the entire grid and conducted replicated (three or four) surveys over two days in each grid in the morning (6:00 hrs. to 10:00 hrs.) and evening (15.30 hrs. to 19.00 hrs.). Detailed methods are available in Khan et. al., (2023). We included this dataset as a reference for a more extensive, focused study to compare our results with.

### Adapting previous surveys for the resurvey

Since the methods used in the past varied, we created a common spatial sampling framework. Worah 1990 did not have GPS locations of the specific points sampled but had detailed maps of sampled forest patches (Supp. Figure S1). We digitized these maps in consultation with Worah. Trivedi 2000 had spatial locations for each transect (Supp. Figure S2), but these were variable in length. Khan 2019 had 45 grids of 1 x 1 km that were sampled. We created 1 x 1 km grids across the landscape to encompass all three sampling strategies. This covered the seven broad locations of the Worah 1990 study, all ten transects of Trivedi 2000, and overlapped with grids from Khan 2019 (including 18 grids where they detected Forest Owlets). Since the transects of Trivedi 2000 encompass multiple adjacent grids, we picked 18 random non-adjacent grids for the re-survey. Together they constituted a total of 46 grids of 1 km x 1km that we selected to resurvey (Figure 1).

### **b)** Are detections of the Forest Owlets at the resurvey locations similar with playback and without?

#### Traditional point count method - without playback

We attempted to recreate the sampling strategy used in Worah 1990 and Trivedi 2000 by visually searching and listening for the call of the Forest Owlet without any playback for 30 minutes within each grid. Two observers AN and AR worked together to cover the entire grid. Before starting this study, both observers were familiar with the species and its calls. This method was carried out before any playback was conducted in the region. Although this method recreates the spatial coverage of the sampled area, it only partially recreates the historic survey method since the re-survey focused on detecting a focal species with a search image. Such a specific search image may also affect the detection of species (Murton 1971; Suzuki 2018).

#### Call playback method

If the Forest Owlet was not detected using the traditional survey method at the same grid, playback of its calls and songs was carried out. The playback procedure followed was the same as Khan 2019, with the same set of calls (AN was part of both field studies). Briefly, playback was conducted at the centre of each grid. If the centroid was inaccessible, a point close to the centre within a 200 m buffer was selected. Playback was conducted for five minutes, followed by an active listening phase of 5 minutes and another final set of playback for five more minutes (a total duration of 15 minutes). At the end of the second playback, the survey was deemed complete if no detection was made earlier. The survey was discontinued for the grid in case the Forest Owlet was detected at any point during this survey period. All calls were played back using a portable Bluetooth speaker, Sony SRS-XB10 connected to a digital recorder (Zoom H1) with a maximum volume of both the speaker and recorder set to match the natural call amplitude of the species (Darras et al. 2018).

All resurveys were carried out between January 2020 and March 2020.

### **c)** Has the landscape or climate changed in the period between the surveys?

#### Detecting Landscape Change between the Survey Periods

We used two methods to detect the landscape change between the historical surveys and the current resurveys. First, we used the Global Forest Change v1.7 (2000-2019) dataset (Hansen et al. 2013). The forest cover loss for the Dang area (with a 5km buffer as described earlier) was extracted in Google Earth Engine (Gorelick et al. 2017). The authors (Hansen et al. 2013) defined Forest Cover Loss as a change from a forest to a non-forest state (from 2000–2019), where the term “forest” refers to tree cover and not land use, and trees were defined as all vegetation taller than 5 meters in height.

Further, we used data from Roy et al. (2015) that mapped the Land-Use and Land-Cover Change in India at decadal intervals for 1985, 1995, and 2005 using the data extracted from multiple imageries and classified following the classification scheme of the International Geosphere-Biosphere Programme (IGBP) (Loveland et al. 2000). We used the Lacos plugin in QGIS (Jung 2013) to calculate the change in land-cover classes in the resurvey grids and across the Dang district, including the 5km buffer zone described previously. Nine landscape classes occurred in the study area of Roy et al. (2015) (details in supplementary methods), and we assessed changes in each of them relevant to the project.

Climate change was examined from 1980 to 2019 across the Dang region within the same 5km. buffer described earlier. We used two types of data to assess these changes.

First, we obtained climate data from Mishra et al. (2019) for the period 01-01-1951 to 31-12- 2019. The resolution of this data is at 0.25 degrees, so we selected the nine grids (Supp. Figure S3) that overlap with the Dang study area (Figure 1). For each grid, we collated data on the mean monthly minimum temperature, mean monthly maximum temperature, and monthly precipitation.

Further, we also assessed change in climate using 19 bioclimatic variables from monthly climate data from Mishra *et al. (2019)* and two other sources for comparison: a) Worldclim 2.5 minutes resolution and an average for the years 1951 - 2000 (“Wordclim Climate Data Version 1.3” 2004) and b) Chelsea remotely sensed bio-climate data of average years 1981 to 2010 at a resolution of 30 arcsec (926 meters) (Karger et al. 2021). We used the biovars function from the *dismo* package (Hijmans et al. 2017) in R. We plotted the spatial changes of select bioclimatic variables across the presence-absence grids that are considered important for Forest Owlet detection, based on the niche models by Koparde et al. (2019) (Supp. Figure S7).

#### Establishment of an acoustic detection framework

a. **d)** Which acoustic analysis software performs better at detecting the calls and songs of the Forest Owlet with co-occurring owls?

#### Data Collection

All acoustic data were collected during the Forest Owlet resurvey study period (January 2020 to March 2020), and all data was recorded between morning 06:00 to 10:00 hrs and evening 14:00 to 18.30 hrs. We collected data with two types of recorders. We used handheld recorders (Zoom H6, F1) to collect several clear target recordings of Forest Owlet vocalizations for training signal recognition software. We also used an Automated Recording Unit (ARU) (Song Meter SM2; Wildlife Acoustics Inc., USA) as a passive recorder, collecting long-term data for testing the Forest Owlet detection in both signal recognition software (described further below).

#### Vocalisations of Forest Owlet

We used two vocalizations of Forest Owlet to test the detector’s efficiency. Additionally, we found a variant of one of the vocalizations while collecting high-quality recordings from handheld recorders in the field. We denote the two main vocalizations as call and song. Call is disyllabic and has a frequency of around 500 to 1100 Hz, (read as ‘kuhuk’..). The song is monosyllabic with a frequency of around 400 to 1400 Hz and longer than the call (read as ‘kwaak…kakak, kwaak…kakak’) (Figure 2).

**Figure 2:**
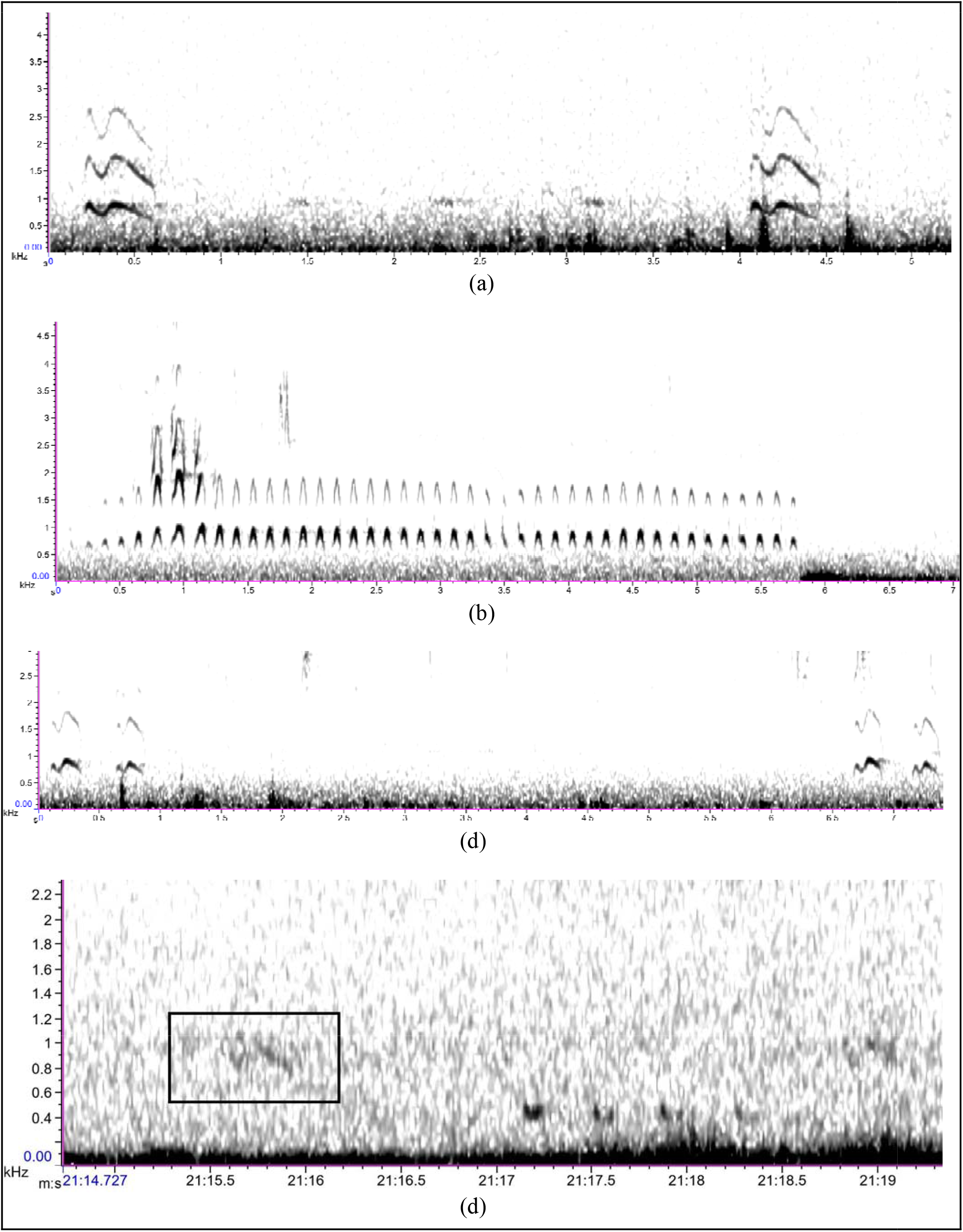
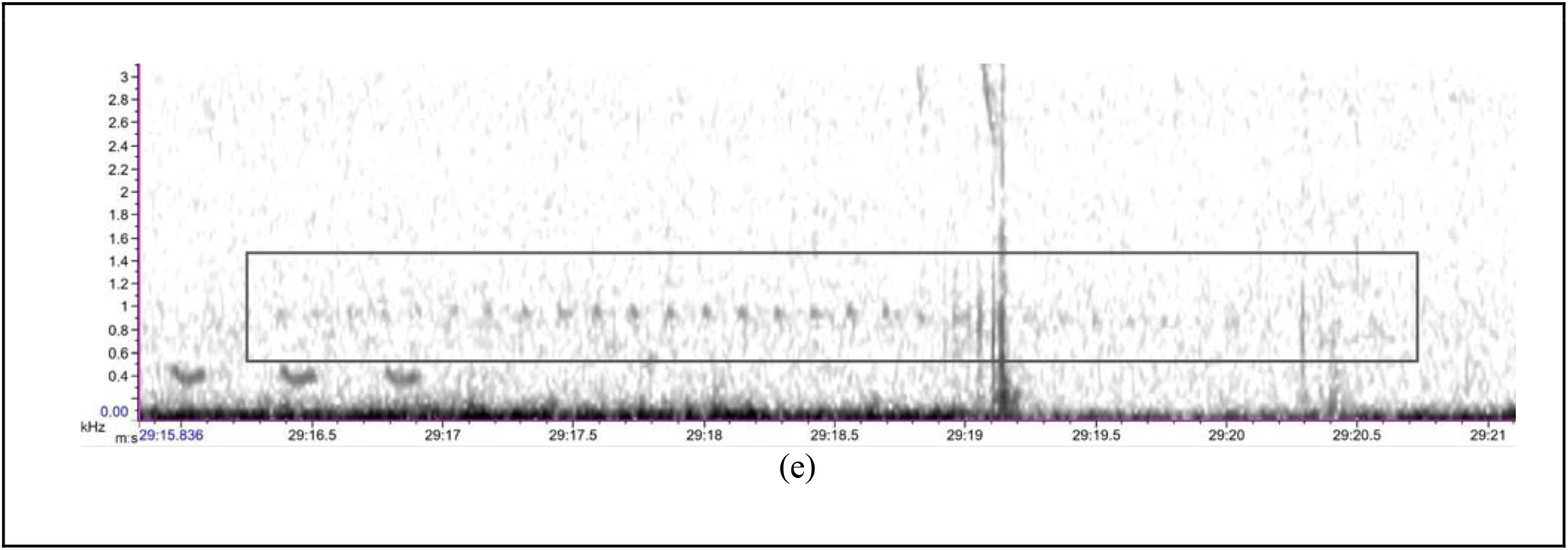
Spectrogram of Forest Owlet vocalizations - a) kuhuk calls, b) kwak kawak songs, c) a variation of kuhuk call, and Spectrogram view of Forest Owlet call (d) and song (e) detection in SM2 recorder at a distance of 300m from the vocalising bird on the field (represented inside the box)

#### Forest Owlet detection using signal detectors

Automated acoustic signal recognition software functions by processing a training template of the target species’ vocalization following the recognition and detection based on the parameters chosen. We selected two signal-detecting software, Raven Pro 2.0 (Bioacoustics Research Program, K. Lisa Yang Center for Conservation Bioacoustics 2018) and Kaleidoscope (“Kaleidoscope Pro Analysis Software 5.6.6” 2024) to create a detector to test the accuracy in detecting the Forest Owlet vocalizations from the field recordings. The recordings collected by the passive recorders (Song Meter SM2; Wildlife Acoustics Inc., USA) during the detection distance estimations were used as a test dataset to run the signal-detecting software for the Forest Owlet detection. A combination of handheld recorder, and ARU data of Forest Owlet vocalisations collected were used as training dataset.

a. Raven Pro 2.0 - Template Detector

Raven Pro 2.0, the acoustic analysis software, requires a good-quality spectrogram template; we obtained these from recordings with handheld recorders (Zomm F1, H6). The template detector function of Raven is used to run the test detection of Forest Owlet vocalization. Effective signal-detecting parameters in Raven were chosen based on the F-Beta values (Arvind et al. 2023; Nolan et al. 2023) by detector validator application. F-beta is the harmonic mean between the precision and recall values of the detector. Based on the F-Beta values, we selected the threshold and frequency values that were better to obtain higher detection rates of Forest Owlet vocalization. The F-Beta for various parameters chosen is calculated by,

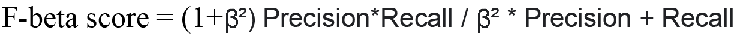

Here, we selected one as the beta to measure the F-beta, which indicates that we gave equal importance to the precision and recall values of the detector performance (Nolan et al. 2023). Based on the results of the F-Beta analysis, we selected specific spectrogram parameters such as window size at 1024, the optimum threshold at 0.65, a frequency of 80 Hz for the Forest Owlet call, and an optimum threshold at 0.45 and a frequency of 100 Hz for the Forest Owlet song to run the detector against the test dataset of 7-hour long recording collected during detection distance experiment using SM2 passive recorders.

### **a)** (b) Kaleidoscope - Advanced Classifiers

The recordings were analyzed by Kaleidoscope Software (version 5.6.6) from Wildlife Acoustics. Kaleidoscope is a signal detection recognizer that builds a classification algorithm by running individual call syllables through Hidden Markov Models (HMM) that maximize the probability of detecting the entire call structure. Kaleidoscope uses K-means clustering of Fisher Scores from a 12-state HMM to cluster all the signals detected into different classes, as opposed to only identifying the signals that match the algorithm above a user-set score threshold (Knight et al. 2017). Advanced cluster analysis was used in the Kaleidoscope detector to reduce the false positive rates since the focus was to detect specific individual species by grouping similar vocalizations in a cluster. This includes a two-step training process and the final testing with the original long-term field recording. The initial two steps involve training the detector by providing good-quality recordings of the Forest Owlet vocalizations from handheld recorders (Zoom H6, F1).

Features used for the training of Kaleidoscope

Kuhuk - Call

500 - 3500 Hz - Freq Range; 0.1 - 0.5 s - Length of detection; 0.35 s - Max inter-syllable gap; 0.5- Max distance; 512 FFT; 12 Max states; 0.5 Max distance to cluster. centre; 30 max clusters

Kwaak - Song

300 - 2000 Hz - Freq Range; 0.1 - 7.5 s- (default) - Length of detection; 0.35 s - Max inter-syllable gap; 1.0 - Max distance; 512 FFT; 12 Max states; 0.5 Max distance to cluster.centre; 50 max clusters

The frequency range was selected to encompass the target calls, and the length of detection and intersyllable and max distance were based on the two types of calls. The number of clusters was set to be larger for kwaak since these are longer songs.

#### Detector performance analysis

The efficiency of both detectors was tested with a common test dataset of a 7-hour recording collected during our field survey. The efficiency of detectors was measured in terms of F-beta score.

#### e) What is the detection distance for Automated Recording Units (ARU) to record the Forest Owlet in its habitat?

We tested the detection distance of the Forest Owlet in the wild. First, we elicited an acoustic response from the Forest Owlet by doing a playback of two vocalisations in the center of the selected grid. Two observers placed ARUs (Song Meter SM2; Wildlife Acoustics Inc., USA) at several distances, viz, 100m, 200m, 300m, 400m, 500m, 600m, and 1000m from the response site of the Forest Owlet in various positive grids. Since it was challenging to have the recorders set out at specific distances from the bird’s calling perch, we selected the 16 positive Forest Owlet locations from our surveys to conduct the distance estimation. Knowing the approximate perch sites allowed us to prepare the ARU placement and only minor adjustments in distance had to be made when the bird responded to our playback. Recording units were programmed to record continuously from the time of deployment with a sample rate at 44100 Hz in Mono using the WAV format with a bit depth of 16 and a gain level between -12 dB to 22 dB.

## RESULTS

In this study, we re-surveyed 46 grids of 1km X 1km that were previously surveyed at different times in the last two decades. We examined the presence of the Forest Owlet with both the traditional and the more recent playback-based techniques.

### Forest Owlets detection at sites that were previously surveyed, differences with playback and without

We detected the Forest Owlet from the areas represented in all three studies, including areas where they were previously not recorded. Additionally, at some of the historic non-detection sites, we could detect Forest Owlet using the traditional visual survey methods (without playback), but the number of detections with playback was considerably higher. We detected the Forest Owlet in 16 grids of the 46 resurveyed. The resurvey of the 18 grids from Khan 2019 with Forest Owlet detections resulted in eight grids with detections in this study (Table 1 and Figure 1). Eight of the 28 grids with no previous Forest Owlet detections (traditional surveys) had Forest Owlet detections during this study.

**Table 1:** Details of resurveys, effort, and detections of the Forest Owlet in the Dang region.

| Surveys by other investigators (Details of Original surveys in Col #1) |  |  | Re-surveys (2020) - this study |  |  |
| --- | --- | --- | --- | --- | --- |
| Team & Year of Survey | No. of grids originally surveyed (A) | Grids (from A) in which Forest Owlet was recorded | No. of grids (from A) re-surveyed (B) | No. of grids Forest Owlet detected during Resurvey (from B) without playback. | No. of grids Forest Owlet detected during Resurvey (from B) with playback |
| Khan 2019 | 45 | 18 | 18 | 2 | 6 |
| Trivedi 2000 | 11 | 0 | 10 | 1 | 1 |
| Worah 1990 | 74 | 0 | 18 | 0 | 6 |

### Landscape Change Analysis

**a) Forest cover change based on Hansen et al.** (2013) **data:** Our assessments of forest cover change over the last two decades across the entire study area of Dang showed an overall minimal decrease of 0.4446 sq. km (Supp. Figure S5), when assessed with global data products (Hansen et al. 2013).
**b) Change in the land cover area based on Roy et al. (2015) data:** In the Dang study area, we detected a 109 sq. km increase in cropland (20% of category) and a decrease in about 46 sq. km of Deciduous Broadleaf Forests (63% of category) and 50 sq.km Fallow land (0.7% of category). Shrubland and Mixed Forest also show a minor decrease in area, and Wasteland shows a minor increase. Although Barren Land and Built-up land were also identified in the study area, no change was detected in this area during this period (Figure 3). This pattern of increase in cropland and a decrease in deciduous forests and fallow land is consistent even when considering a larger landscape (5 km buffer around Dang) (Supp. Figure S6).

More specifically, in the 46 resurvey grids where the Forest Owlet was not recorded had an increase in Cropland, Deciduous Broadleaf Forest, and Water Bodies, and a decrease in Shrubland and Mixed Forest between 1985 and 2005. Meanwhile, the grids where the Forest Owlet was recorded are characterized by a slight increase in Cropland and Mixed Forest and a decrease in Deciduous Broadleaf Forest and Water Bodies in the same period (Figure 3).

**Figure 3:**
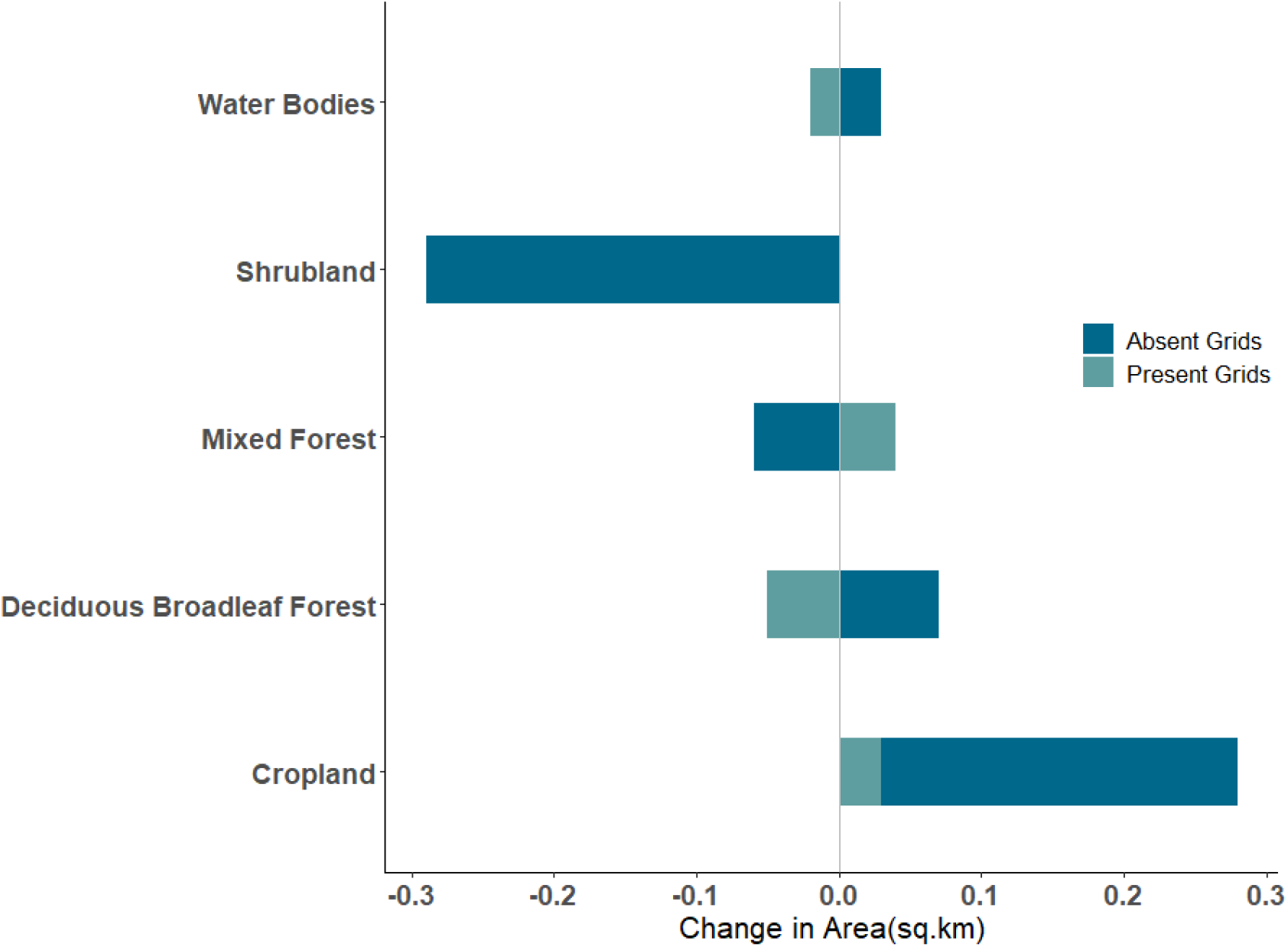
Change in the area (sq. km) of land use types from 1985-2005 in resurvey (presence and absence of Forest Owlet) grids of Dang based on data from Roy et al. (2015) (classes described in detail in Supplementary Materials)

### Climate change analysis

Based on the datasets we examined, we did not recover any major differences in annual mean minimum temperature over the years, but there was a significant difference in the annual max temperature in 2019 compared to 1980 and 2000 (Figure 4).

**Figure 4:**
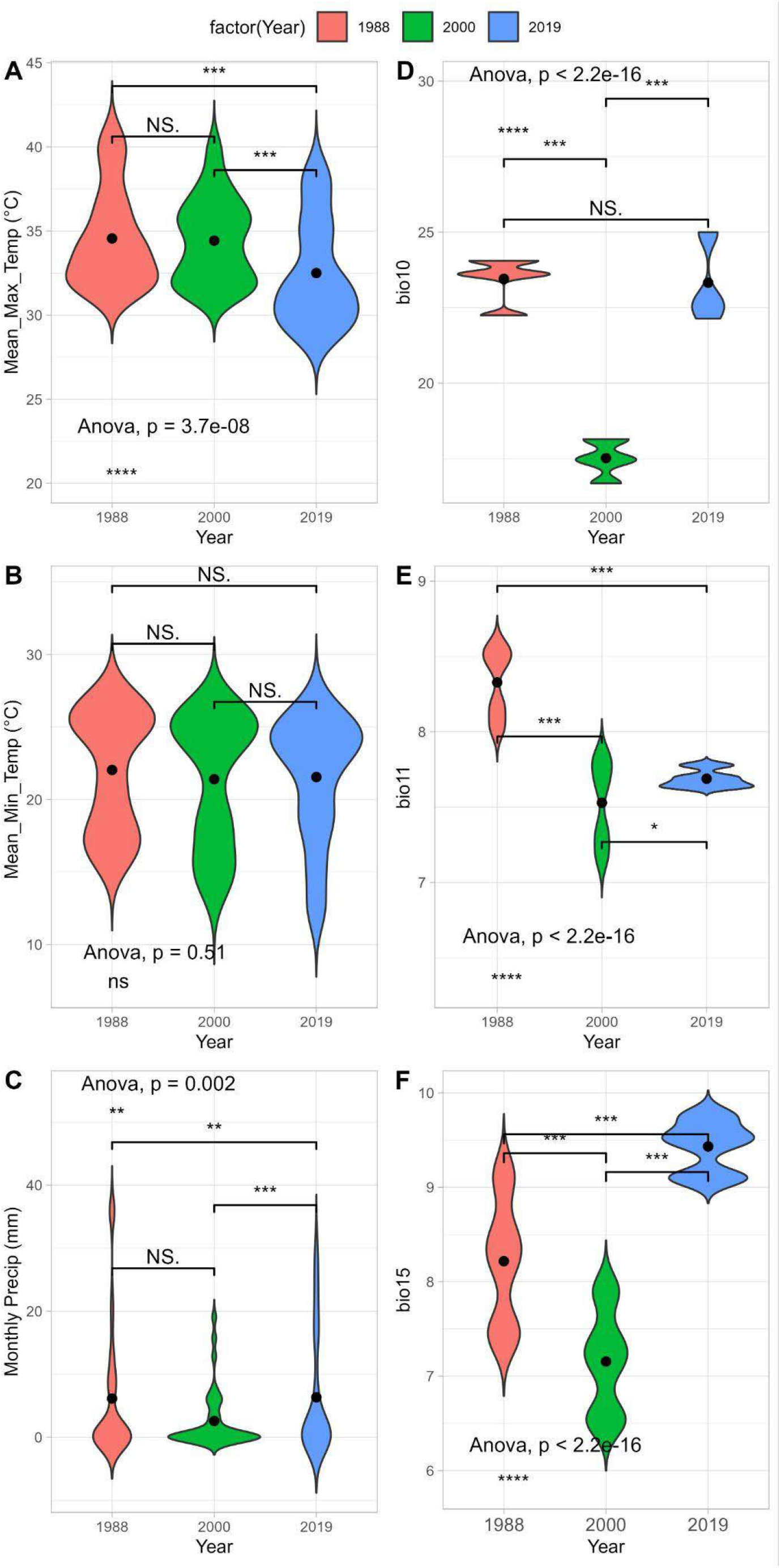
A- Annual mean maximum temperature (°C), B - Annual mean minimum temperature °C), C - Monthly mean Precipitation in mm, D - Bio 10, Mean temperature of the warmest quarter, E - Bio11, Mean temperature of the coldest quarter, F - Bio15, Precipitation Seasonality (Coefficient of Variation) across Dang (5 km buffer - 9 climate points) * indicates significance level (* P ≤ 0.05, ** P ≤ 0.01, *** P ≤ 0.001, **** P ≤ 0.0001 (For the last two choices only) and ns - non-significant, Black dot indicates the mean values

### Establishment of an acoustic detection framework

#### Detector performance analysisilo06p;l6y

A total of 124 calls and 83 songs of the Forest Owlet vocalization were detected from the 7 hours of field recording using automated recorders. We found that the Kaleidoscope detector performed better at detecting the shorter Forest Owlet call with a higher precision of 0.7, recall of 0.5, and F-beta score of 0.6 (Figure 5). However, the Forest Owlet song was detected better in the Raven detector with a relatively higher precision of 0.5, recall of 0.24, and an F-beta score of 0.31 (Figure 5).

**Figure 5:**
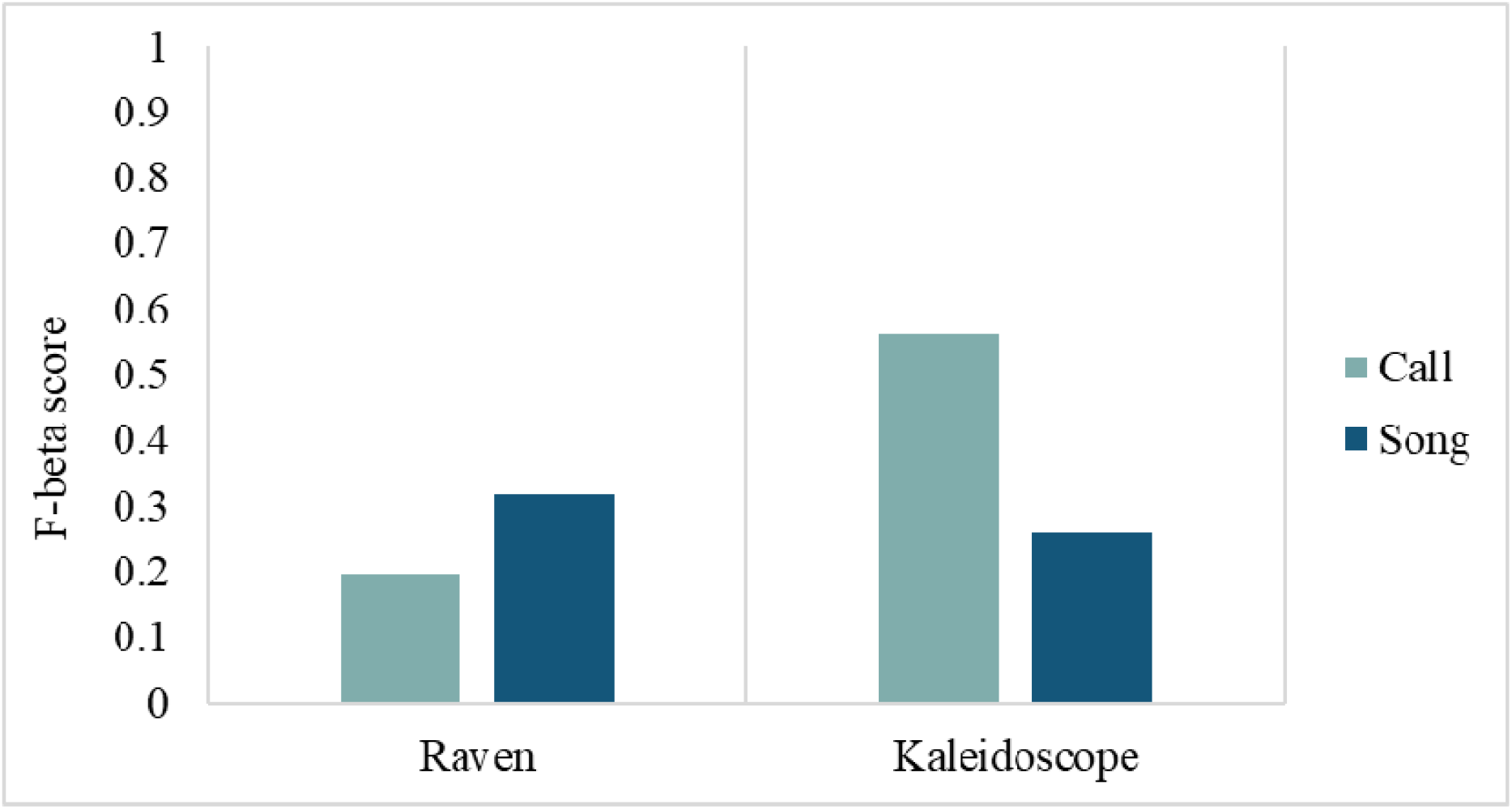
F-Beta score of Raven and Kaleidoscope in the detection of Forest Owlet vocalizations

#### Detection distance analysis

By placing SM2 recorders from 100 to 1000 meters at various intervals, we could detect the clear Forest Owlet acoustic vocalizations at a distance of up to 300 meters from the real bird acoustic activity (Figure 2).

## DISCUSSION

In this study, to assess the presence of the Forest Owlet, we resurveyed 46 sites (1x1 km grids) previously surveyed by multiple researchers. We detected the Forest Owlet in areas where previous studies had not reported them.

We consider three possibilities with varying survey methods - search image, playback, and temporal replicates, and two possible biological reasons - landscape change and climate change for this outcome.

### Role of a Search Image

Numerous studies have indicated that studies that are not actively searching for a target species, such as in surveys of broad communities, often miss cryptic species (Dawkins 1971; Lauriault and Wiersma 2019). In India, broad studies of Western Ghat birds surmised that the skulking rainforest bird, *Sholicola* was rare (Daniels et al. 1991; Kannan 1998) while targeted surveys found it to be common at high elevations (Robin and Sukumar 2002). Here, two datasets (Worah 1991) and (Trivedi 2006) were broad bird community studies from the landscape and were conducted with no search image of this target species at a time when the Forest Owlet was not known to occur in the landscape. Our detection of the Forest Owlet from the exact locations does throw up the possibility that the bird was present but not detected.

### Role of playback

Several bird detection studies have documented that the use of playback (of calls or songs) increased the detection probability relative to the passive surveys (Haug and Andrew B. Didiuk 1993; Celis-Murillo, Deppe, and Allen 2009; Boscolo, Metzger, and Vielliard 2006). The detection of owls is more effective with call playbacks (Haug and Andrew B. Didiuk 1993; Conway and Simon 2003). The male burrowing owls *Speotyto cunicularia* were very responsive and immediately responded to the playback by singing back the primary song, bobbing, and flying toward the source (Haug and Andrew B. Didiuk 1993). Although we find increased detection and responses to playback, the behavioral ecology of this response is not very clear. The first information on the Forest Owlet vocalizations and behaviour was provided by (Rasmussen and Ishtiaq 1999), and only more recent surveys (Ishtiaq and Rahmani 2004; Mehta et al. 2008; Khan et al. 2023), were able to use this method. We note that most numbers of reports of the Forest Owlet have increased since the publication of its vocalizations.

### Significance of temporal replicates

We find that even with playback, there is variation in the detection of this rare species. Khan et al. (2023) note that the detection probability of the species is ∼70% in Dang with multiple temporal replicates. Of the 18 grids where they detected Forest Owlet in Dang, 13 (72%) grids had positive detections in all four of their survey replicates. However, just a year later, we did not detect them in all of these grids. Significantly, our study did not include temporal replicates, and some non-detection may be associated with detection probability. Most importantly, although there is stochasticity in the detection (or presence) of this relatively rare bird, even when the first temporal replicate of Khan et al. (2023) is considered, 18 of 45 grids were positive, which is similar to our detections (8 of 18 grids). This pattern holds even if we consider the identity of the grid. Within the design constraints of the current study, we cannot ascertain if this stochasticity is due to the probability of detection or a biological process such as local movement. This is where we suggest acoustic monitoring, described further below, comes in.

### Relationship with climate and landscape change data

Numerous studies across the globe have found species occurrences responding to both climate and landscape changes. Resurveys of Iknayan and Beissinger (Iknayan and Beissinger 2018) in a desert landscape indicate that climate change may be related to the change in bird communities over the decades. However, in some cases, species like understory insectivorous birds are more vulnerable to land-use changes (Sreekar et al. 2015). Our study finds some changes over the decades with the raw climate data, and also with the derived/combined climate data. Some of these climatic variables were shown to be relevant for broader Forest Owlet distribution based on a model created by Koparde et al. (2019). However, both outputs need to be considered with the caveat of the spatial resolution and scale of the original data. Original climate data of (Mishra et al. 2019), Worldclim, and Chelsea are at very large scales (details in the methods section) and may not capture changes that occur at the scale of our study area. There are few ground sensors in this region to examine such patterns for this period. Our study did not find any change in landscape to forest cover as measured with global products (Hansen et al. 2013). However, we do find changes based on a regional data product (Roy et al. 2015), and the changes are consistent with an increase in agriculture and a reduction in deciduous forests and fallow lands (indicative of open habitats). We note that field observations by Worah 1990 and Trivedi 2000 suggest that agriculture has increased, and the teak plantations have matured and become larger. Such details are not reflected in the current dataset and may well impact local site preferences. The type of agricultural practice - retaining old-growth trees, may also have facilitated the presence of the Forest Owlet. We also cannot rule out associated biological processes like competition with co-occurring owls - Jungle Owlet and Spotted Owlet, interacting with the habitat and climate change. Much of these remain to be examined.

From our assessments of the various detectors, the longer vocalizations of the Forest Owlet - the songs are better detected with RAVEN, while the shorter calls are detected with Kaleidoscope. This is consistent with the findings that each species may vary in the detection based on specific signatures (Scott Brandes 2008). In this case, the two types of vocalisations of a species are detected differently (Romero-Mujalli et al. 2021). Our assessment of the use of bioacoustics as a survey method included an assessment of the detection distance of the bird’s call in its habitat. We found that for calls and songs that are clear enough to be detected with RAVEN or Kaleidoscope, the recorders must be placed 300-400m from the vocalizing Forest Owlet. We note that this distance is different from human detection distance (up to ∼1km (Ishtiaq and Rahmani 2004)), as the current detection distance is specifically for clear spectrograms that can be detected with these algorithms. This distance estimate is best suited for the deciduous forests of Dang; in more open landscapes, the distance may be larger. Based on the extensive surveys of Khan et al. (2023), we are unsure that the species may be found in denser forests. Hence, this distance (300m radius) could be used as a conservative grid size for placing ARUs for long-term acoustic monitoring.

### Recommended long-term monitoring techniques

a. Considering the stochasticity/variability in the detection of the Forest Owlet, we recommend using temporal replicates to detect the species.
b. The use of playback has increased very rapidly in recent efforts in bird detection surveys, and our study suggests that playback is useful for the detection of the Forest Owlet. However, several authors (Watson, Znidersic, and Craig 2019; Harris and Haskell 2013), indicate that excessive playback can cause abandonment of territories, higher levels of anxiety (Budka et al. 2019), and may impact other crucial behavioral activities (Harris and Haskell 2013). In other cases, some birds may not respond similarly to playback (de Lima and Roper 2009), and some may stop responding to excessive playback (Jepson 2011). Although our study did not explicitly test the impacts of playback, we do not recommend durations longer than 5 minutes at an amplitude louder than the natural calls.
c. We recommend long-term monitoring of the Forest Owlet using Automated Recording Units. These can also be deployed at likely Forest Owlet sites to understand the species’ range better. We recommend a gird size of 600m (based on a 300m detection distance from this study) in deciduous forests.

In summary, we find Forest Owlet in locations where they were not previously detected, and we find a positive impact of using playback for surveys. We, however, do not attempt to make any causal links between the decadal detections of the endangered Forest Owlet and climate or landscape change. Although we note the changes indicated in climate and landscape over this duration, we suggest detailed monitoring of climate and landscape in these and surrounding areas for future monitoring. Based on the current study, we suggest survey protocols for the long-term monitoring of the Forest Owlet.

## ACKNOWLEDGEMENTS

We thank MOEFCC (Grant No:J.22012/61/2009-CS(W), dated: 29th September 2017) to SM, VVR, Rajah Jayapal and IISER Tirupati institutional funding to VVR; Gujarat Forest Department, particularly Shri Agneeshwar Vyas, DCF- Dang South Forest Division for permits, suggestions to reach survey locations, and beat guard officers for accompanying few days in the field; AR and AN thanks PG and his wife Sangeetha for their hospitality, Rawal and Shivshakti for the help with field travel logistics; Several people assisted with analyses - Jobin Varughese, Arasumani M with GIS; Vijay Ramesh - curating climate data from IIT-Guwahati; Viral Joshi, Chiti Arvind - for help with acoustics ideas and analyses. PG thanks the GEER Foundation, Gandhinagar (Gujarat) for coordinating and funding the study. VVR thanks Pradyumna Gogte for alerting us of the presence of historical data from the landscape - the catalyst for initiating this study. We also thank the bird lab group at IISER Tirupati including Ashwin Warudkar, Vishwa Jagati, Isha Bopardikar for their constructive comments and feedback on the manuscript.

## AUTHOR CONTRIBUTIONS

Conceptualisation, Resources: VVR (lead) and SM, Data Curation and Investigation: AR, AN

Formal analyses, and Visualisation: AR (lead), MRM Funding: VVR and SM - equal

Supervision: VVR (equal lead), SM (equal lead), PG, SW Project administration: VVR and SM

Writing - Original Draft: VVR (lead), AR, AN, MRM

Writing - Review & Editing - VVR, AR (lead), AN, SM, PG, SW

---

**Figure S1:**
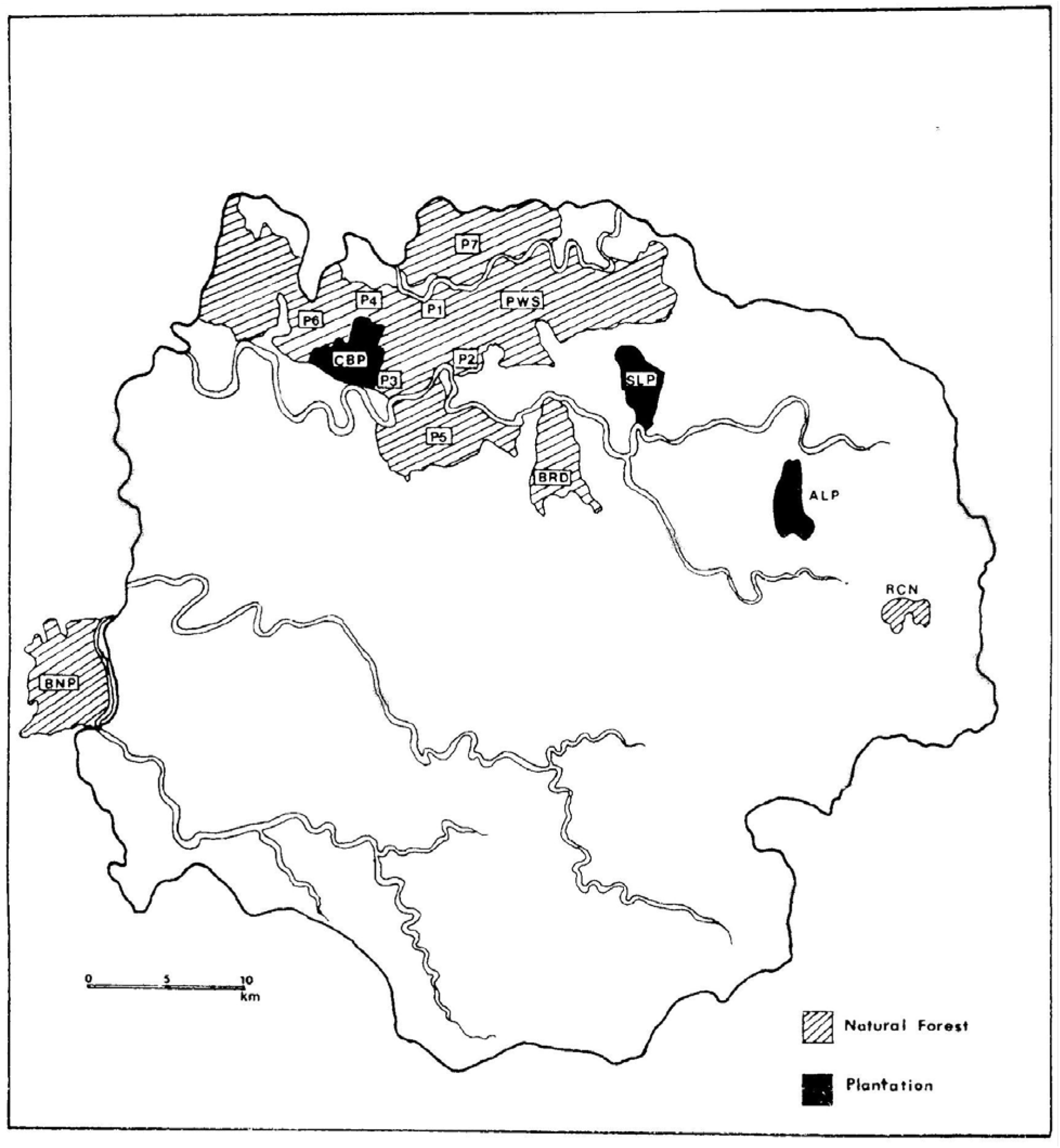
Sampled areas of Worah 1990 study area across Dang

**Figure S2:**
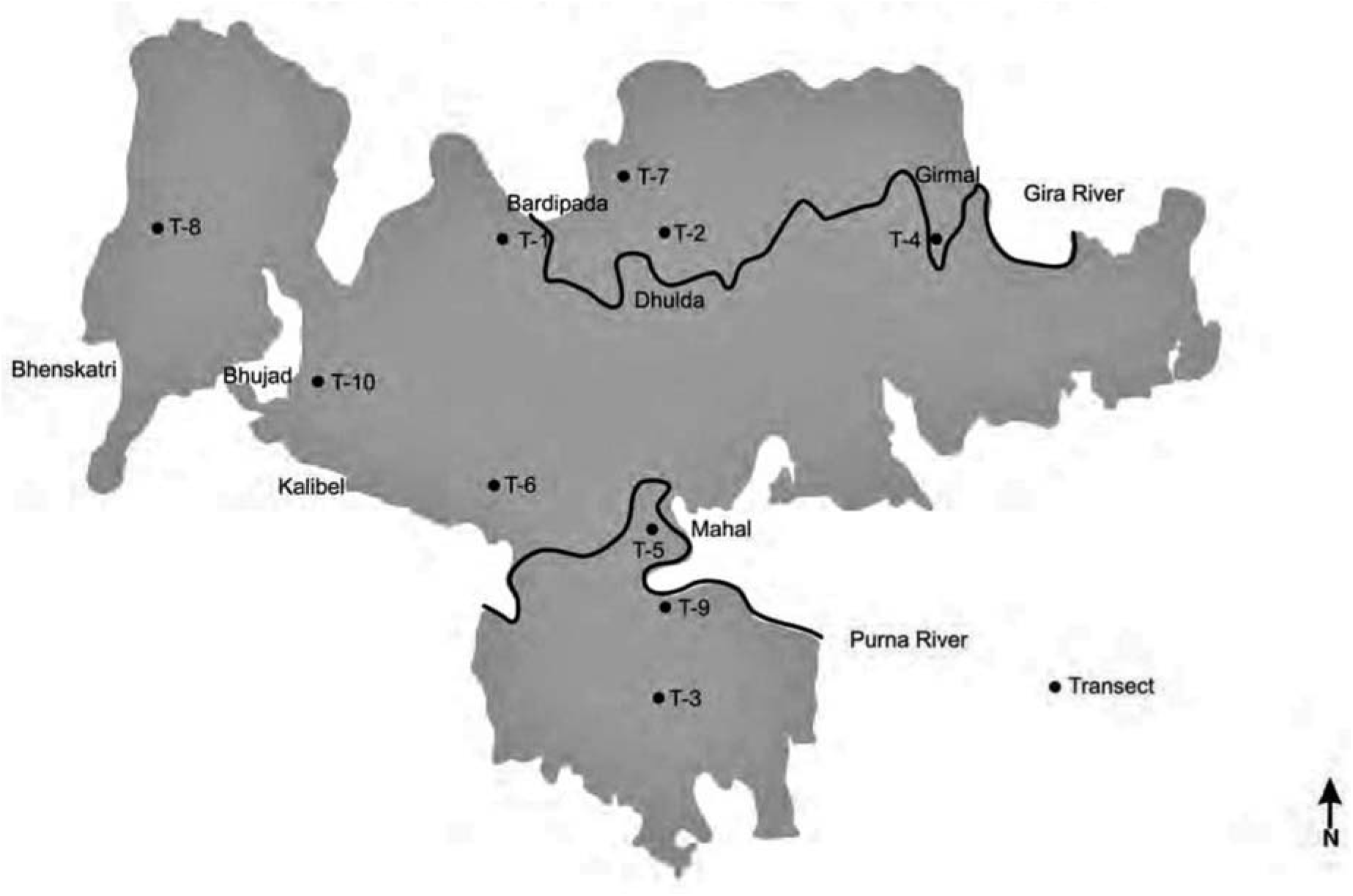
Locations of transects by Trivedi 2000 across Dang, including Purna Wildlife Sanctuary, north of Dang

**Figure S3:**
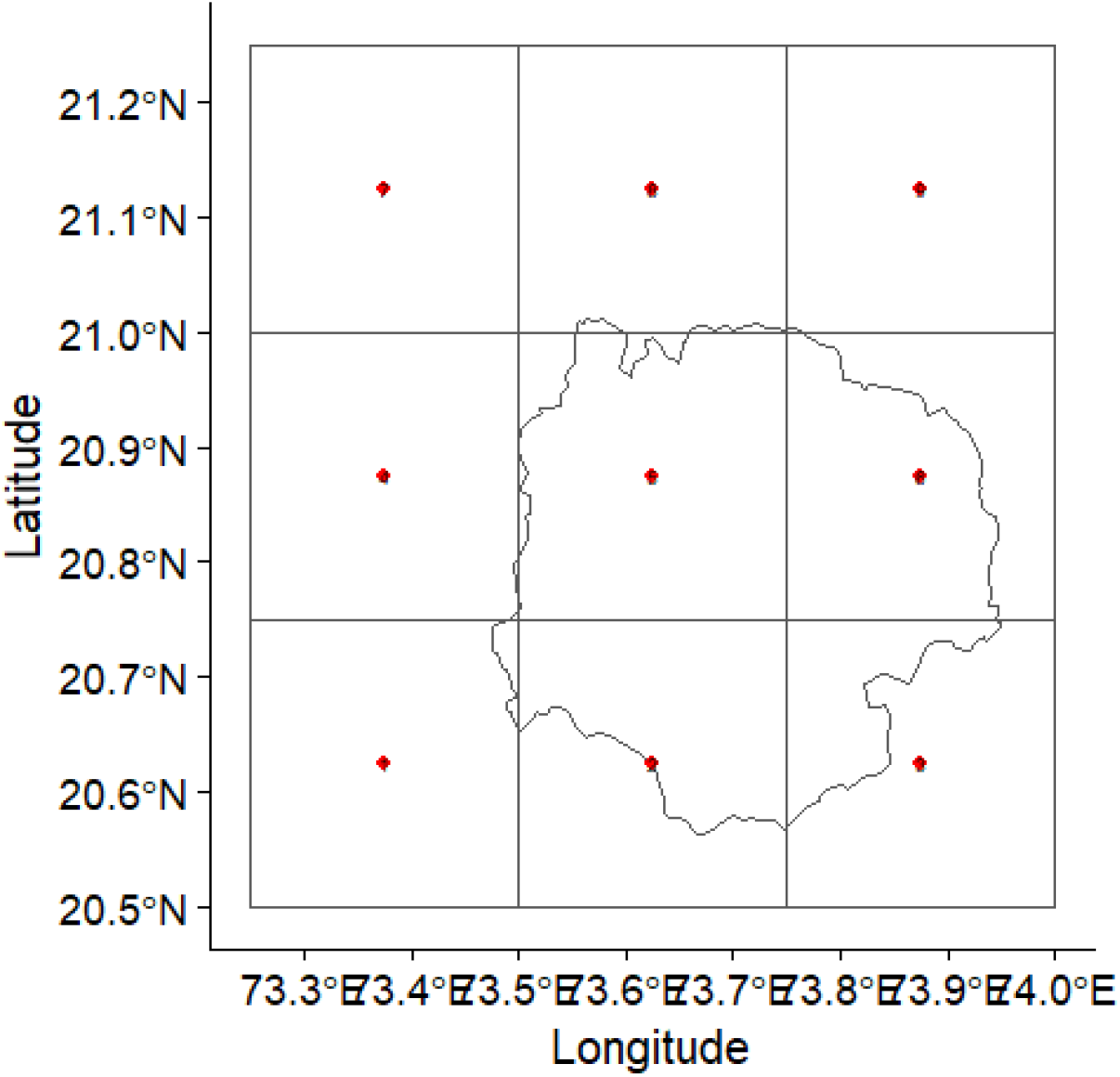
Locations/centroids of 0.25-degree grids around the Dang region (outline map) for which historic climate data from 1951 to 2019 based on Mishra *et al. (2019)* was extracted for comparison with present data

**Figure S4a:**
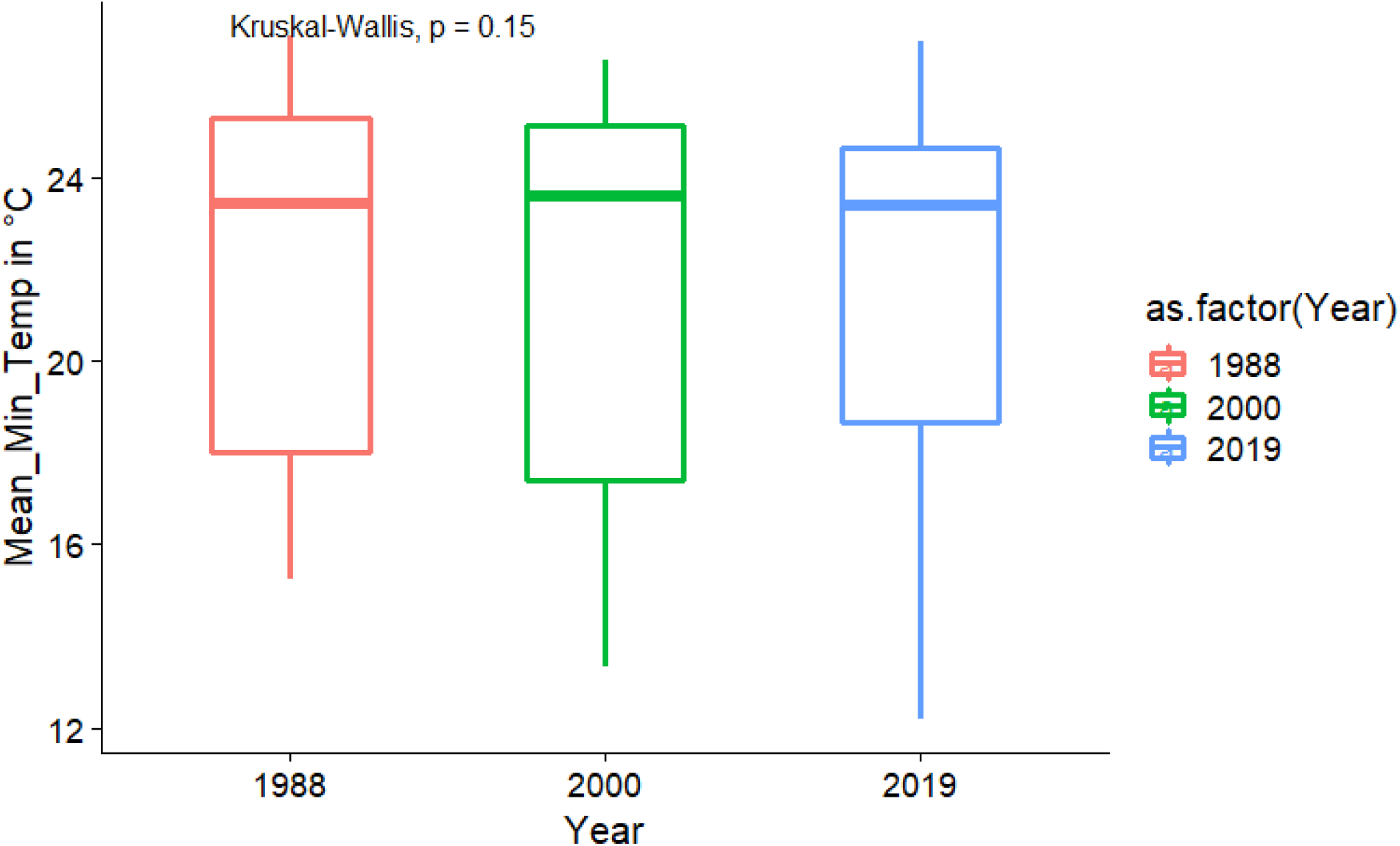
Annual mean minimum temperature based on Mishra *et al. (2019)* across Dang over three time periods closest to the survey years shows no significant change

**Figure S4b:**
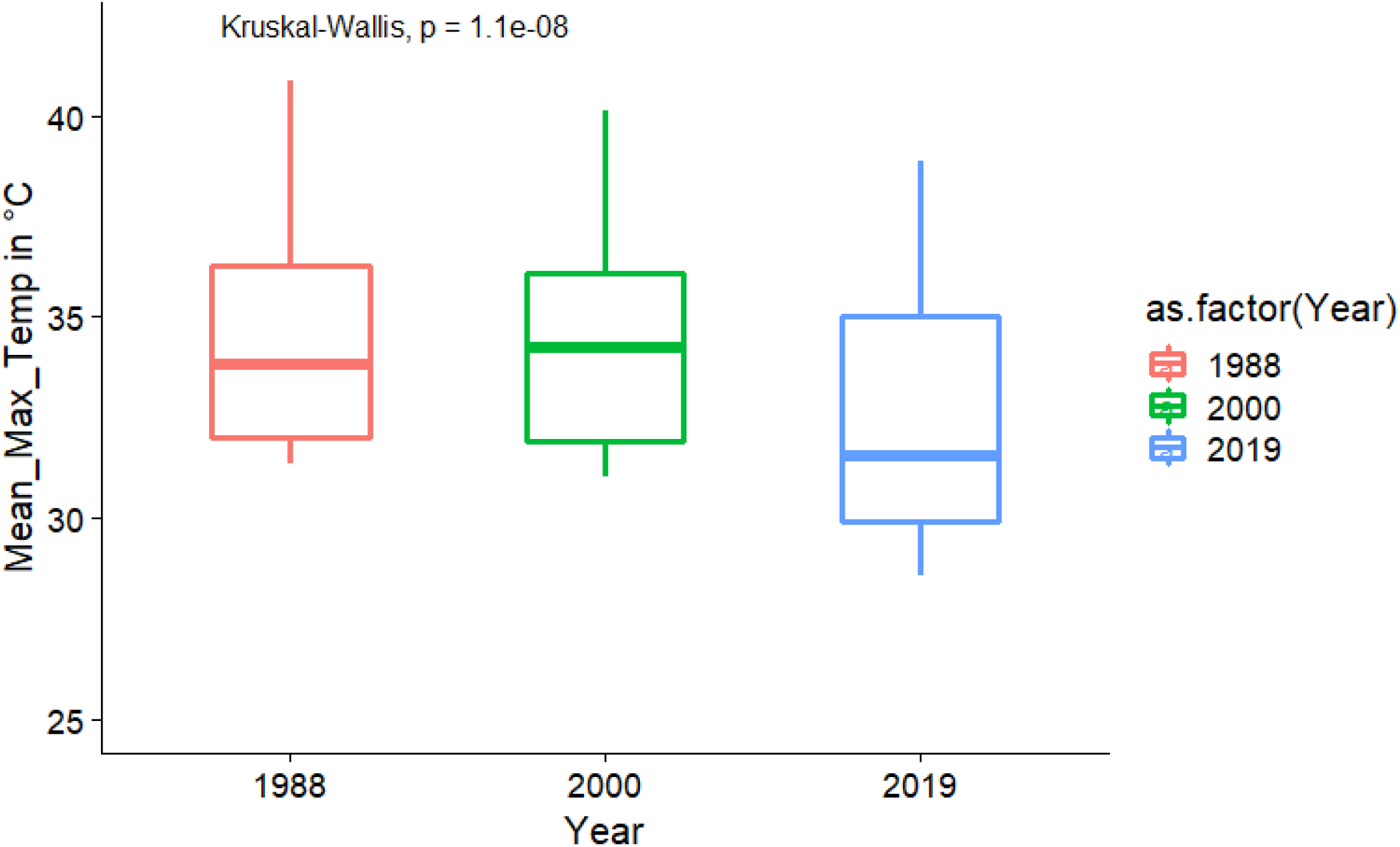
Annual mean maximum temperature based on Mishra *et al. (2019)* across Dang over three time periods closest to the survey years shows a significant change in 2019

**Figure S4c:**
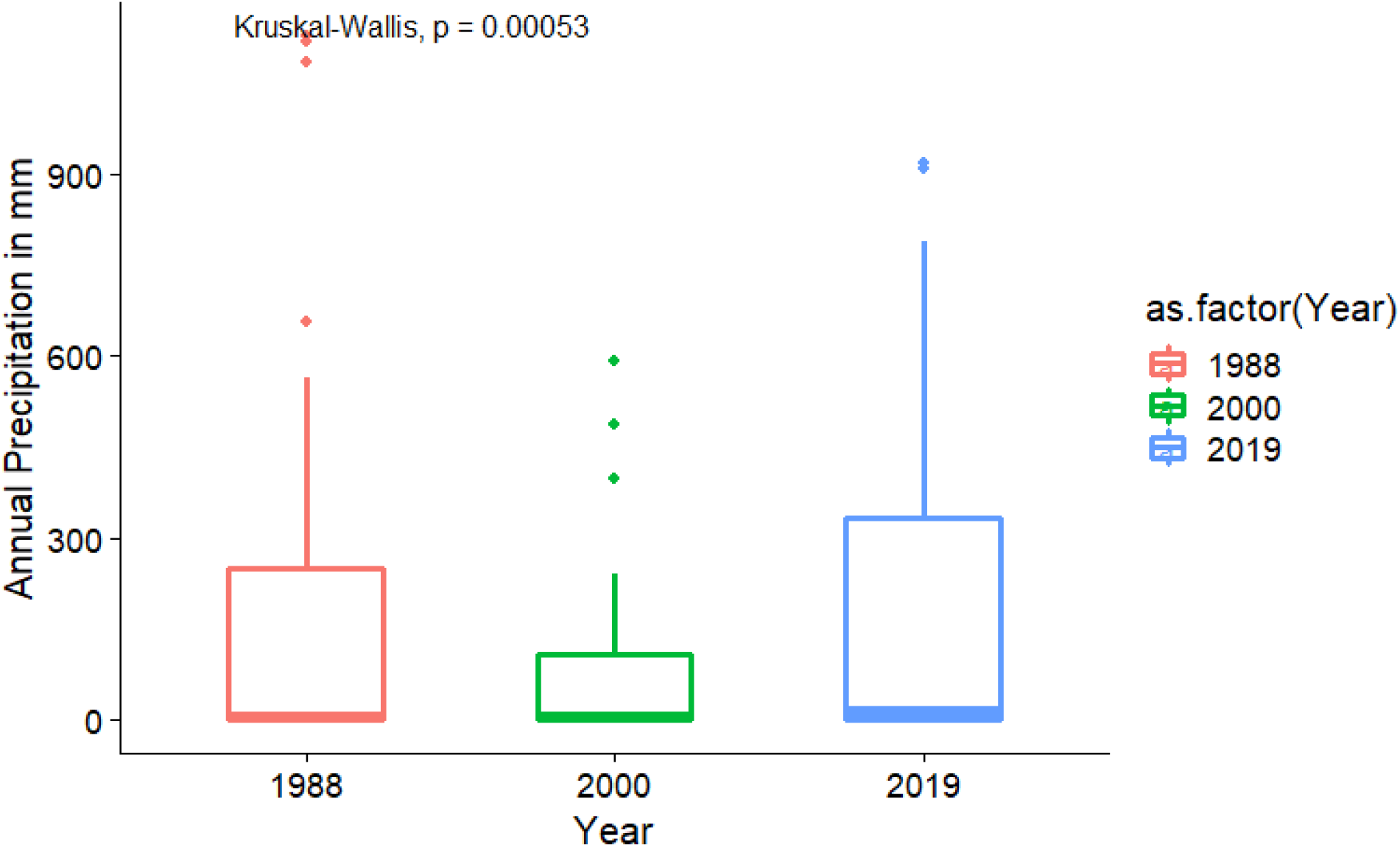
Mean annual precipitation based on Mishra et al. (2019) across Dang over three time periods closest to the survey years shows no significant change

**Figure S5:**
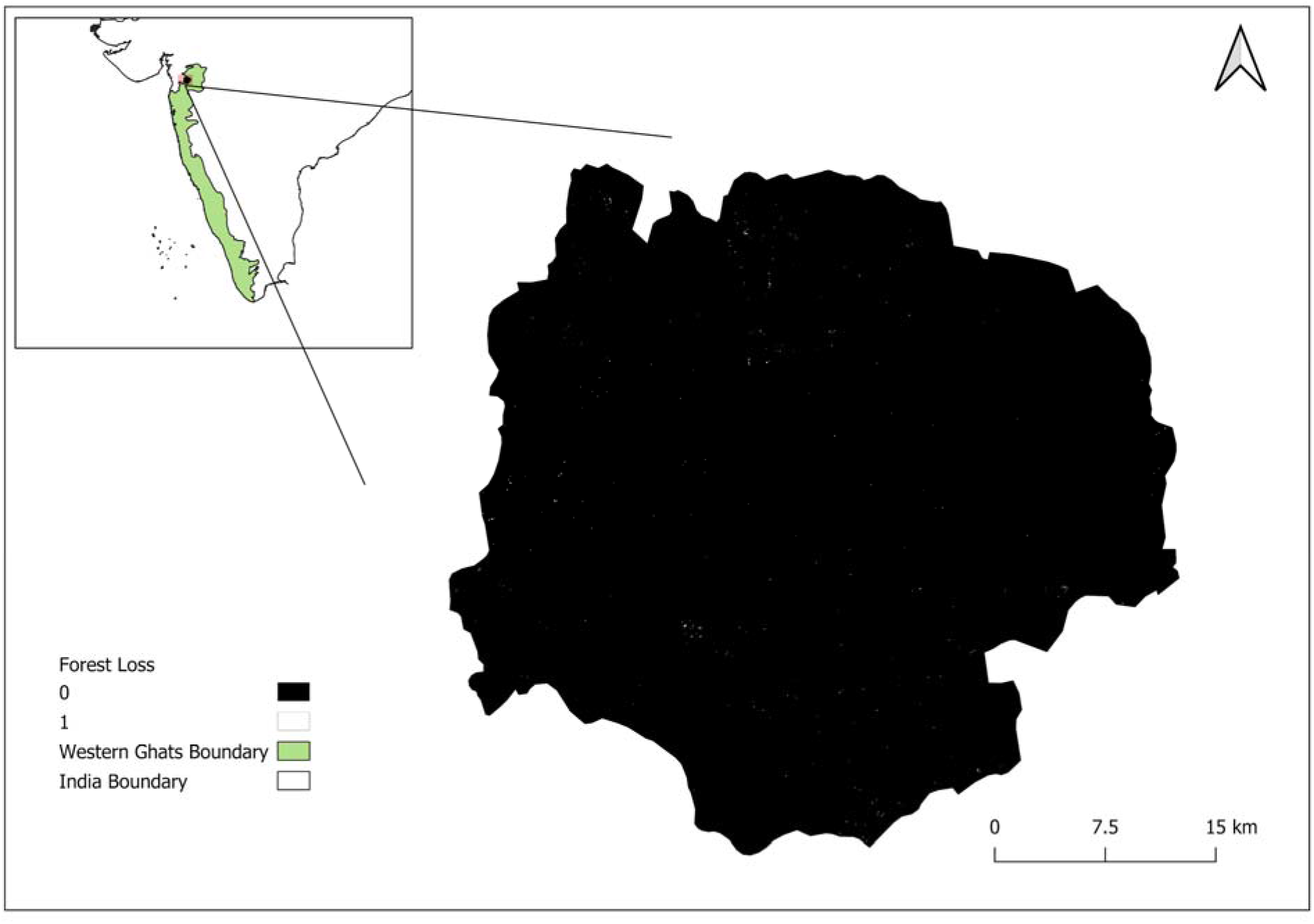
Forest loss across Dang from 2000 to 2019 appears minimal when assessed with global data products (Source: Hansen/UMD/Google/USGS/NASA). Here, pixel value 1 represents forest loss; 0 represents no perceivable change.

**Figure S6:**
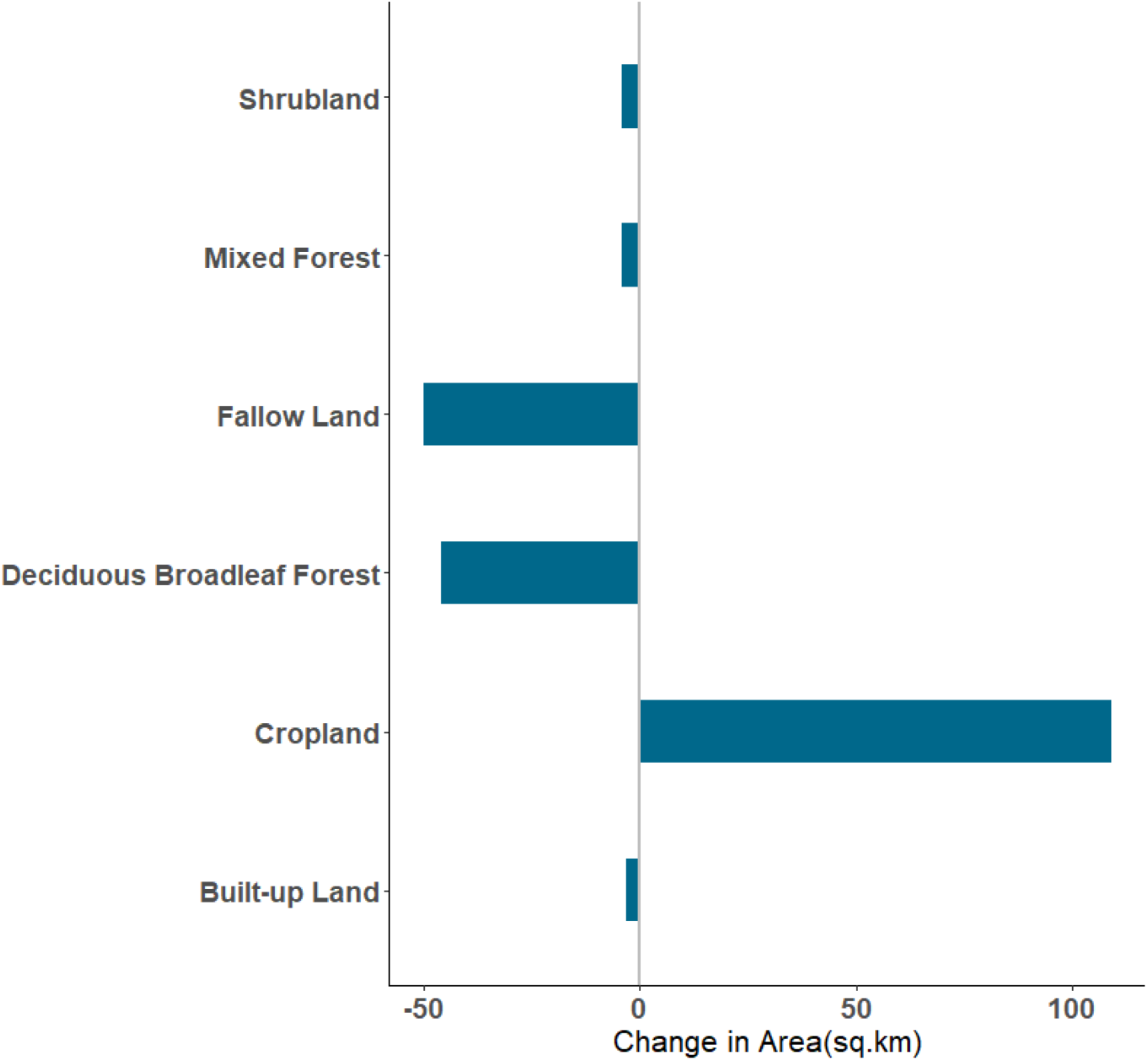
Change in area of land use types from 1985-2005 in the Dang district based on Roy *et al. (2015)*, with a 5km buffer - indicating landscape change at a larger landscape scale

**Figure S6:**
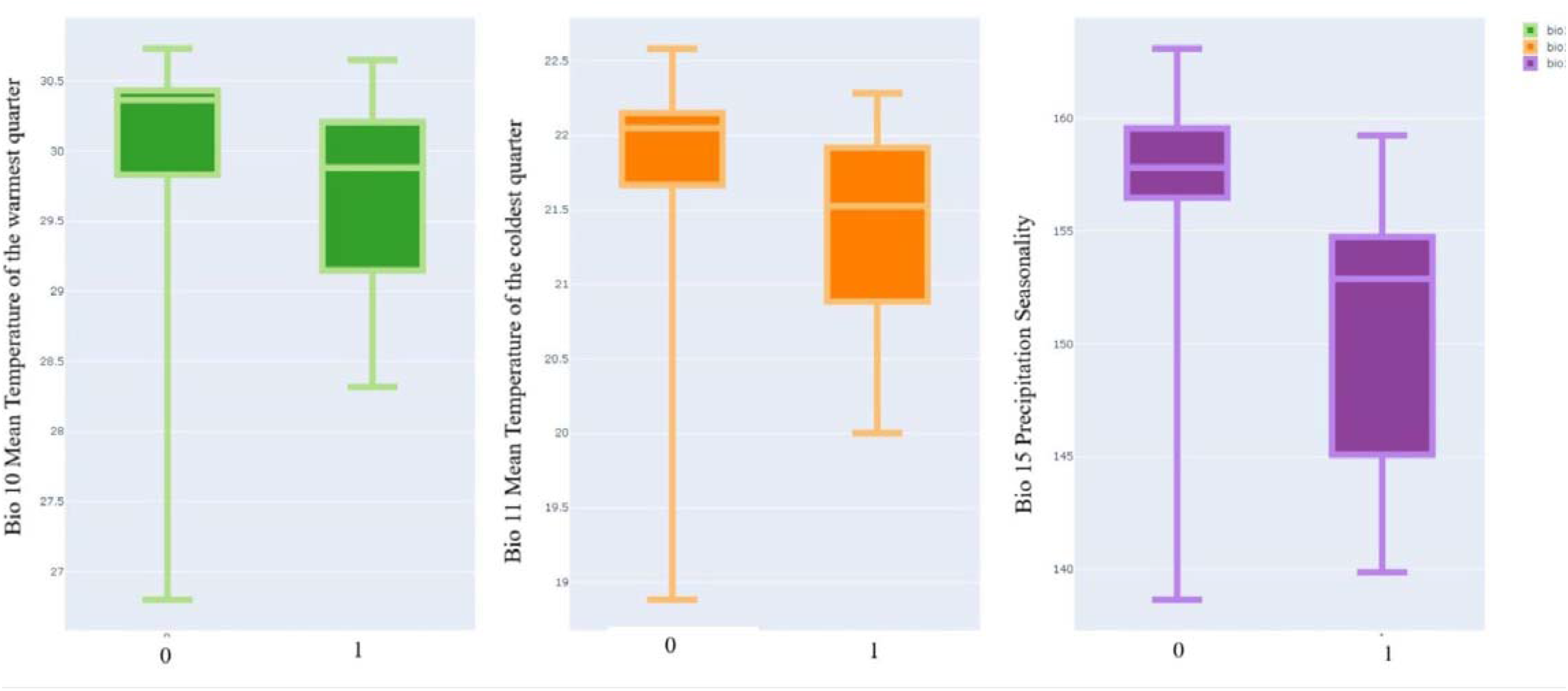
Selected bioclimatic variables based on Worldclim Climate data version 1.3 (“Wordclim Climate Data Version 1.3” 2004) at locations with Forest Owlet detections across Dang (1 - presence, 0 - absence)

**Figure S7:**
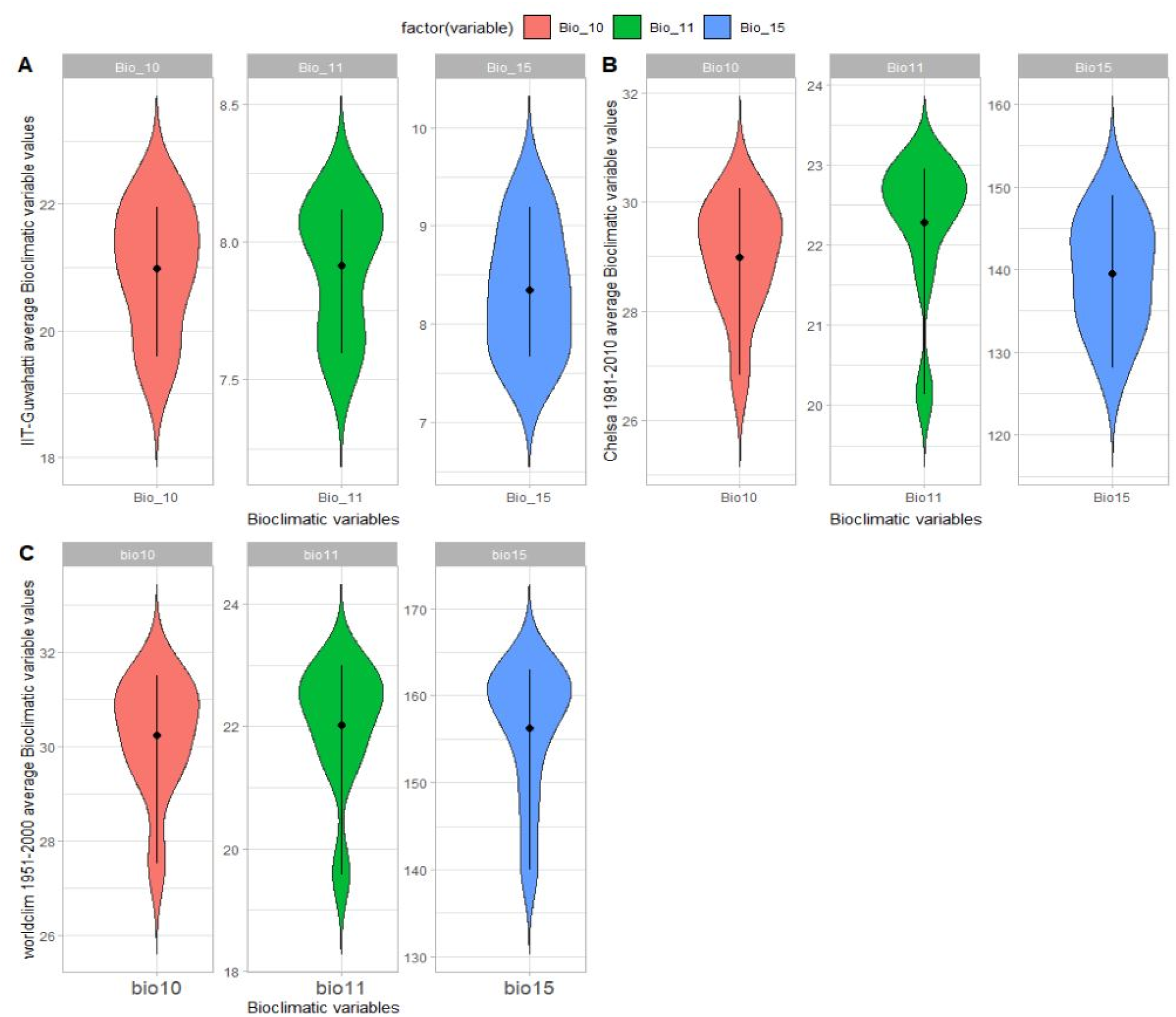
Comparison of selected bio-climatic variable values based on Chelsea (Karger et al. 2021) (average of years 1981 to 2010) and Worldclim Climate data version 1.3 (“Wordclim Climate Data Version 1.3” 2004)(average of years 1951 to 2000) across nine grids of climate locations from Mishra et al. (2019) climate data

## SUPPLEMENTARY MATERIALS

## List of supplementary materials

1. Supplementary methods - land-cover classes from Roy et al. (2015),
2. Figure S1: Sampled areas of Worah 1990 study area across Dang
3. Figure S2: Locations of transects by Trivedi 2000 across Dang, including Purna Wildlife Sanctuary, North of Dang
4. Figure S3: Locations/centroids of 0.25-degree grids for which historic climate data was extracted for comparison with present data
5. Change in Climate

a. Figure S4a: Annual mean minimum temperature across Dang over three time periods closest to the survey years shows no significant change
b. Figure S4b: Annual mean maximum temperature across Dang over three time periods closest to the survey years shows a significant change in 2019
c. Figure S4c: Mean annual precipitation across Dang over three time periods closest to the survey years shows no significant change
6. Figure S5: Forest loss across Dang from 2000 to 2019 appears minimal when assessed with global data products (Source: Hansen/UMD/Google/USGS/NASA)
7. Figure S6: Selected worldclim bioclimatic variables at locations with Forest Owlet detections across Dang.
8. Figure S7: Chelsea (average of years 1981 to 2010) and Worldclim (average of years 1951 to 2000) selected the bio-climatic variable across 9 grids of climate locations from IIT-Guwahati climate data

## Supplementary methods

The land-cover classes identified in the study area by Roy *et al*. (2015) are:

1. **Deciduous Broadleaf Forest** - Woody vegetation with a percent cover >60% and height exceeding 2 m. Consists of broadleaf tree communities with an annual cycle of leaf-on and leaf-off periods.
2. **Cropland** - Temporarily cropped area followed by harvest and a bare soil period (e.g. single and multiple cropping systems). Note that perennial woody crops will be classified as either forest or shrubland, whichever is appropriate. Includes orchards. Different types of cropland based on seasons (e.g. kharif, rabi, zaid) were not subclassified.
3. **Builtup Land** - Land covered by buildings and other man-made structures
4. **Mixed Forest** - Trees with a percent cover >60% and height exceeding 2 m. Consists of tree communities with interspersed mixtures or mosaics of the other four forest types. None of the forest types exceeds 60% of landscape
5. **Shrubland** - Land with woody vegetation less than 2 m in height and with greater than 10% shrub canopy cover. The shrub foliage can be either evergreen or deciduous
6. **Barren Land** - Exposed soil, sand, rocks, or snow and never have more than 10% vegetated cover during any time of the year
7. **Fallow Land** - Land taken up for cultivation temporarily allowed to remain uncultivated for one or more seasons.
8. **Waste Land** - Sparsely vegetated land with signs of erosion and land deformation that could be attributed to lack of appropriate water and soil management, or natural causes. These are land identified as currently underutilized and could be reclaimed to productive uses with reasonable effort. Degraded forest (<10% tree cover) with signs of erosion is classified under wasteland
9. **Water Bodies** - Areas with surface water, either impounded in the form of ponds, lakes, reservoirs or flowing as streams, rivers, etc. Can be either fresh or salt- water bodies.

